# The piRNA cluster *torimochi* is an expanding transposon in cultured silkworm cells

**DOI:** 10.1101/2022.09.07.506900

**Authors:** Keisuke Shoji, Yusuke Umemura, Yukihide Tomari

## Abstract

PIWI proteins and PIWI-interacting RNAs (piRNAs) play a central role in repressing transposable elements in animal germ cells. It is thought that piRNAs are mainly produced from discrete genomic loci named piRNA clusters, which often contain many “dead” transposon remnants from past invasions and have heterochromatic features. In the genome of silkworm ovary-derived cultured cells called BmN4, a well-established model for piRNA research, *torimochi* was previously annotated as a unique and specialized genomic region that can capture transgenes and produce new piRNAs bearing a trans-silencing activity. However, the sequence identity of *torimochi* has remained elusive. Here, we carefully characterized *torimochi* by utilizing the updated silkworm genome sequence and the long-read sequencer MinION. We found that *torimochi* is in fact a full-length gypsy-like LTR retrotransposon, which is exceptionally active and has massively expanded its copy number in BmN4 cells. Many copies of *torimochi* in BmN4 cells have features of open chromatin and the ability to produce piRNAs. Therefore, *torimochi* may represent a young, growing piRNA cluster, which is still “alive” and active in transposition yet capable of trapping other transposable elements to produce *de novo* piRNAs. (185 words)

## INTRODUCTION

Transposons are DNA sequences that can migrate from one region to another in the genome. Since transposon insertions can disrupt the genetic structure, it is necessary for the hosts to suppress transposon activity especially in germ cells, where the genome is inherited to the next generation (Ernst, Odom and Kutter, 2017). PIWI-interacting RNAs (piRNAs) and PIWI proteins play a central role in suppressing transposons in the germline (Klattenhoff and Theurkauf 2008; Ghildiyal and Zamore 2009). PIWI proteins use piRNAs as the sequence-specific guide to repress target transposons in two ways: transcriptional gene silencing (TGS) and post-transcriptional gene silencing (PTGS). piRNA-mediated TGS is achieved by heterochromatic histone modifications and DNA methylation (Alexei A Aravin et al., 2008; Sienski, Dönertas and Brennecke, 2012). On the other hand, piRNA-mediated PTGS relies on the endoribonucleolytic activity of PIWI proteins (Li et al. 2009a; Reuter et al. 2011; Wang et al. 2014; de Fazio et al. 2011). The 3′ fragments of the cleavage products by piRNA-guided PIWI protein are incorporated into another PIWI protein and processed into new piRNAs (Brennecke et al. 2007; Gunawardane et al. 2007). These new piRNAs further guide the cleavage of complementary RNAs (i.e., the same strand as the original piRNA), leading to the production of piRNAs with the same sequence as the original piRNAs. This cleavage-dependent piRNA biogenesis, called the “ping-pong” amplification cycle, is thought to be broadly conserved among many animals including flies, mice, zebrafish, sponges, and silkworms (Brennecke et al., 2007; Houwing et al., 2007; Alexei A. Aravin et al., 2008; Grimson et al., 2008; Kawaoka et al., 2009).

Most piRNA sequences are densely mapped to discrete genomic regions, called piRNA clusters (Brennecke et al. 2007). The piRNA clusters often contain a variety of fragmented, “dead” transposon remnants deriving from past invasions, producing piRNAs that can recognize and repress currently active transposons (Gunawardane et al. 2007; Nishida et al. 2007; Saito et al. 2006; Reuter et al. 2011; de Fazio et al. 2011). In this way, piRNA clusters are thought to act as genomic storage devices that keep the information of non-self sequences to be suppressed.

The *flamenco* and *42AB* loci in *Drosophila melanogaster* represent the most well-studied piRNA clusters (Brennecke et al. 2007). Both piRNA clusters are more than 100 kbp long and possess trimethylation of Lys9 on histone H3 (H3K9me3), a typical mark of heterochromatin (Rangan et al. 2011). These large heterochromatic piRNA clusters were originally proposed to provide the immunity to homologous transposon copies across the genome *in trans* (Malone and Hannon, 2009; Levine and Malik, 2011). However, it was recently shown that these large piRNA clusters are evolutionarily labile and mostly dispensable for transposon suppression, raising the possibility that dispersed elements in individual transposons can mediate silencing of themselves *in cis* (Gebert et al., 2021). Moreover, it has been reported that new euchromatic transposition of transposons can act as a source of piRNAs (Shpiz et al. 2014; Olovnikov et al. 2013). Such euchromatic piRNA clusters are found not only in flies but also in mice and mosquitos (Alexei A. Aravin et al., 2008; Shpiz et al., 2014; George et al., 2015). These findings suggest that piRNA clusters are not stationary entities but can flexibly appear and disappear.

BmN4 cells, derived from silkworm ovaries, retain the active piRNA pathway including the ping-pong amplification cycle, and thus have been widely used in piRNA research (Kawaoka et al. 2009). A previous study using BmN4 cells identified *torimochi* (the Japanese word for birdlime) as a piRNA cluster in silkworms (Kawaoka et al. 2012).

*Torimochi* was regarded as a unique and specialized piRNA cluster in the BmN4 genome, which traps foreign transgenes and produces piRNAs that can repress homologous transgenes in other genomic regions *in trans*. However, the detailed sequence properties of *torimochi* have remained unclear.

The previous *torimochi* study was conducted using the old silkworm genome sequence originally published in 2008 (The International Silkworm Genome Consortium 2008; Kawaoka et al. 2012). At that time, the silkworm genome contained many unassembled regions, even within *torimochi*. However, the silkworm genome sequence has recently been updated with completing most of the previously unassembled regions (Kawamoto et al. 2019). In this study, we carefully reexamined the *torimochi* region to understand why it can trap foreign transgenes for silencing. Using the updated silkworm genome sequence and the long-read sequencer MinION, we found that *torimochi* is in fact a full-length gypsy-type transposon, which has massively expanded its copy number in BmN4 cells compared to silkworms. Moreover, our single-nucleotide polymorphism (SNP) and chromatin immunoprecipitation (ChIP) analyses indicated that many copies of *torimochi* in BmN4 have features of open chromatin and the ability to produce piRNAs. Therefore, *torimochi* may serve as a model for young piRNA clusters, which are still “alive” and active in transposition, can trap other transposons, and produce *de novo* piRNAs.

## RESULTS

### The piRNA cluster *torimochi* is a full-length gypsy-type transposon

In the original silkworm genome sequence published in 2008, *torimochi* was annotated as a specialized piRNA cluster located on chromosome 11, because no highly homologous region was found elsewhere in the genome (Kawaoka et al. 2012; The International Silkworm Genome Consortium 2008). Moreover, the chromosome 11 *torimochi* itself contained a large unassembled region. However, recent improvements in sequencing technology have determined the sequences of many previously unassembled regions in the silkworm genome (Kawamoto et al. 2019). Using the updated silkworm genome sequence published in 2019, we performed a nucleotide BLAST search using the known *torimochi* sequence as a query. We found that there are at least three copies of *torimochi*-like sequences, not only on chromosome 11 but also on chromosomes 12 and 24 (Fig. 1A). Alignment of these three copies of *torimochi*-like sequences suggested that they are full-length gypsy-type retrotransposons, which even retain the conserved LTR sequences at both ends (Fig. 1B). Therefore, *torimochi* is not a unique sequence in the genome but should be now interpreted as a gypsy-type transposon.

**Figure 1.**
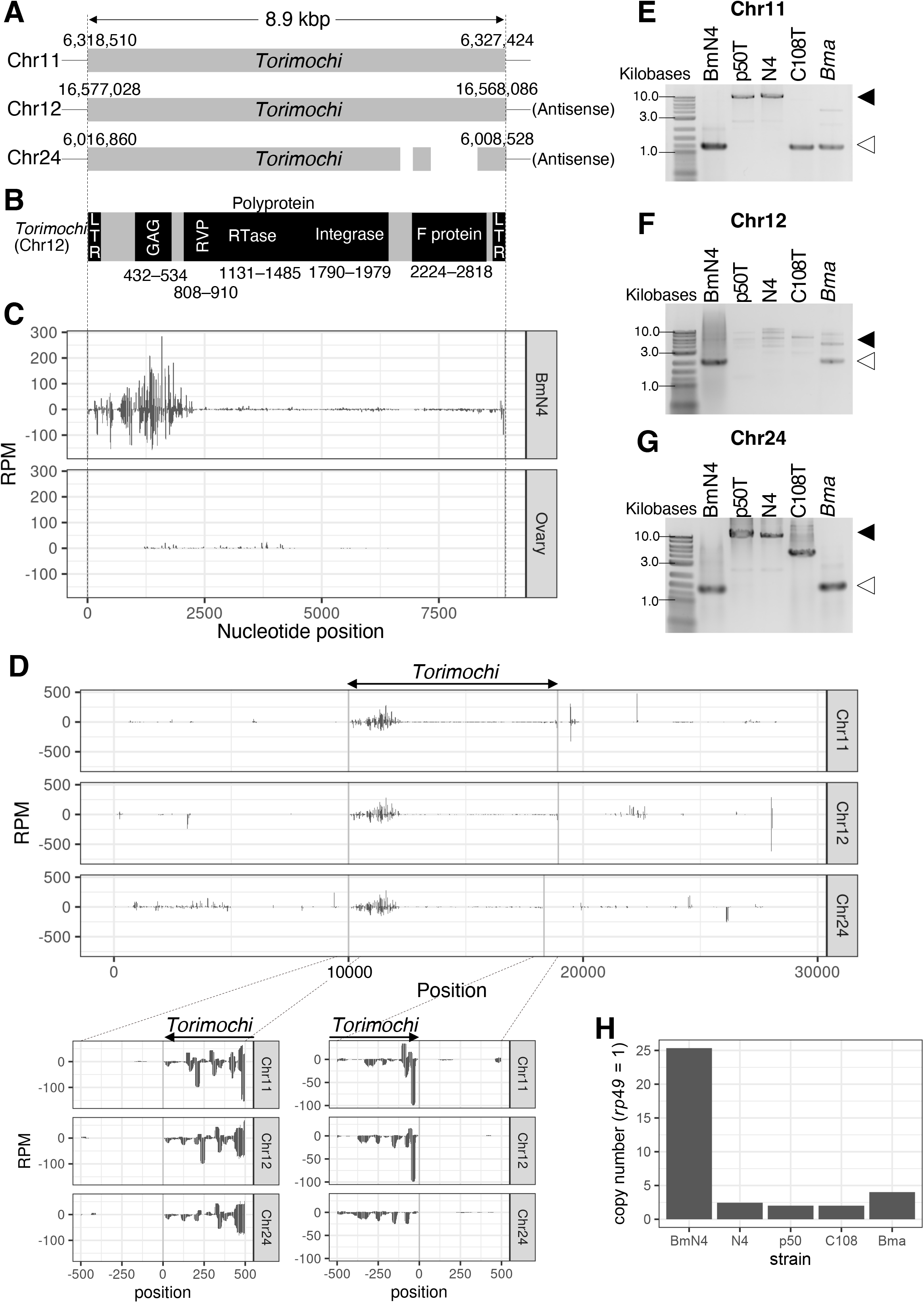
*torimochi* is a full-length, gypsy-type transposon. (A) Schematic representation of the three copies of *torimochi* in the recently published silkworm genome. The copy on chromosome 24 has a deletion in the region encoding F protein. (B) Structure of *torimochi* on chromosome 12. There are LTR sequences at both ends, with retrotransposon gag protein (GAG), retroviral aspartyl protease (RVP), reverse transcriptase (RTase), integrase, and F protein (Baculovirus F protein) encoded inside. RVP, RTase, and integrase are encoded as polyproteins without stop codons. Black boxes show the reading frame and the numbers below indicate the amino acid numbers. (C) Distribution of *torimochi*-derived piRNAs in BmN4 cells and silkworm ovaries. Although *torimochi* on chromosome 11 is used as a representative, piRNAs produced from *torimochi* on other chromosomes can be mapped in the same way due to sequence similarity. The positive and negative values on the y-axis indicate the coverage of piRNAs in the sense and antisense directions, respectively (the same hereafter). (D) Distribution of piRNAs derived from the regions inside and outside of *torimochi* in the recently published silkworm genome. (E–G) Genomic PCR using primers outside of *torimochi* on chromosome 11 (E), chromosome 12 (F), and chromosome 24 (G). The genome DNAs of BmN4 cells and silkworm strains p50T, N4, and C108T and *B. mandarina* (*Bma*) were used. Black, white arrows indicate the band lengths with and without *torimochi*, respectively. (H) Estimation of the copy number of *torimochi* by qPCR of genomic DNA. The copy number of *rp49* (a gene on an autosomal chromosome) was normalized to 1.

We found that *torimochi*-derived piRNAs are highly expressed in BmN4 cells but not in silkworm ovaries, suggesting that *torimochi* has gained the piRNA-producing ability in BmN4 cells (Fig. 1C). To further characterize the piRNA-generating activity of *torimochi*, we compared the patterns of piRNA production inside and outside of *torimochi* for the three insertion sites on the BmN4 genome. Every insertion site showed clear boundaries for piRNA production; piRNAs were densely mapped within the *torimochi* region, whereas the outside region was almost devoid of piRNAs (Fig. 1D). This observation suggests that *torimochi* is a full-length transposon that acts as a piRNA-producing cassette.

The 2019 silkworm genome was based on the p50T strain of silkworm (Kawamoto et al. 2019). Given that *torimochi* is a full-length transposon that could in theory retain transposition activity, the actual locations of *torimochi* may be different between the genomes of P50T strain and BmN4 cells. Therefore, we performed quantitative genomic PCR of three silkworm strains (p50T, N4, and C108T), *Bombyx mandarina* (*Bma*, a presumed ancestor of *Bombyx mori*), and BmN4 cells. We found that the *torimochi* insertion status at the above-identified three genomic locations greatly varies among species. In fact, the BmN4 genome lacks *torimochi* insertion at any of the three insertions (Fig. 1E–G). Nevertheless, when we measured the copy number of *torimochi* by qPCR, we found that *torimochi* in BmN4 cells has ∼25-fold more copies than the r*ibosomal protein 49* (*rp49*) gene on the autosomes, which is remarkably higher than those in the *B. mori* strains and *B. mandarina* (Fig. 1H). These results suggest that *torimochi* is an active transposon unit that keeps transpositioning in the silkworm genome and that BmN4 cells have gained massive copy numbers of *torimochi* presumably during the establishment of the cell line and/or cell passage.

### Detailed annotation of *torimochi* in the BmN4 genome by utilizing MinION sequencer

To investigate where *torimochi* copies are inserted in the BmN4 genome, we opted to use the long-read sequencer MinION. We processed the MinION reads to search for the junctions between the known *torimochi* sequences and other genomic regions in BmN4 cells (Fig. 2A). Among them, we identified at least nine sites that have reliable junctions at both ends and with a reasonable genomic distance. Genomic PCR showed two distinct lengths of bands—with and without *torimochi*—for eight of those nine sites, indicating that *torimochi* is inserted in a heterozygous manner at each site (Fig. S1). For the insertion in chromosome 22, bands of incomplete lengths were observed, so we decided to exclude it from further analyses for simplicity (Fig. S1). We then obtained the precise sequences of these eight *torimochi* insertions by Sanger sequencing of PCR-amplified fragments and constructed a phylogenetic tree together with the above-identified three *torimochi* insertion sequences in the p50T silkworm genome. We found that all the eight *torimochi* sequences in the BmN4 genome are similar to each other, belonging to one clade that is distant from the three *torimochi* copies in p50T strain (Fig. 2B). This suggests that expansion of *torimochi* in BmN4 cells has occurred relatively recently. In the eight homologous *torimochi* sequences in the BmN4 genome, we could identify 61 SNP positions that can help distinguish them (Fig. 2C). At each position of these SNPs, we analyzed the nucleotide composition of BmN4 piRNAs using previously published data (Izumi et al. 2020). We found piRNA sequence polymorphisms at all the SNP positions (Fig. 2D, E), suggesting that *torimochi* piRNAs are produced from multiple *torimochi* loci, rather than a single locus. Comparison of the piRNA production patterns inside and outside of *torimochi* for these eight insertion sites showed clear boundaries for piRNA production (Fig. S2A–H). These observations support the model that *torimochi* is a full-length, multi-copy transposon, each copy of which acts as a piRNA-producing cassette.

**Figure 2.**
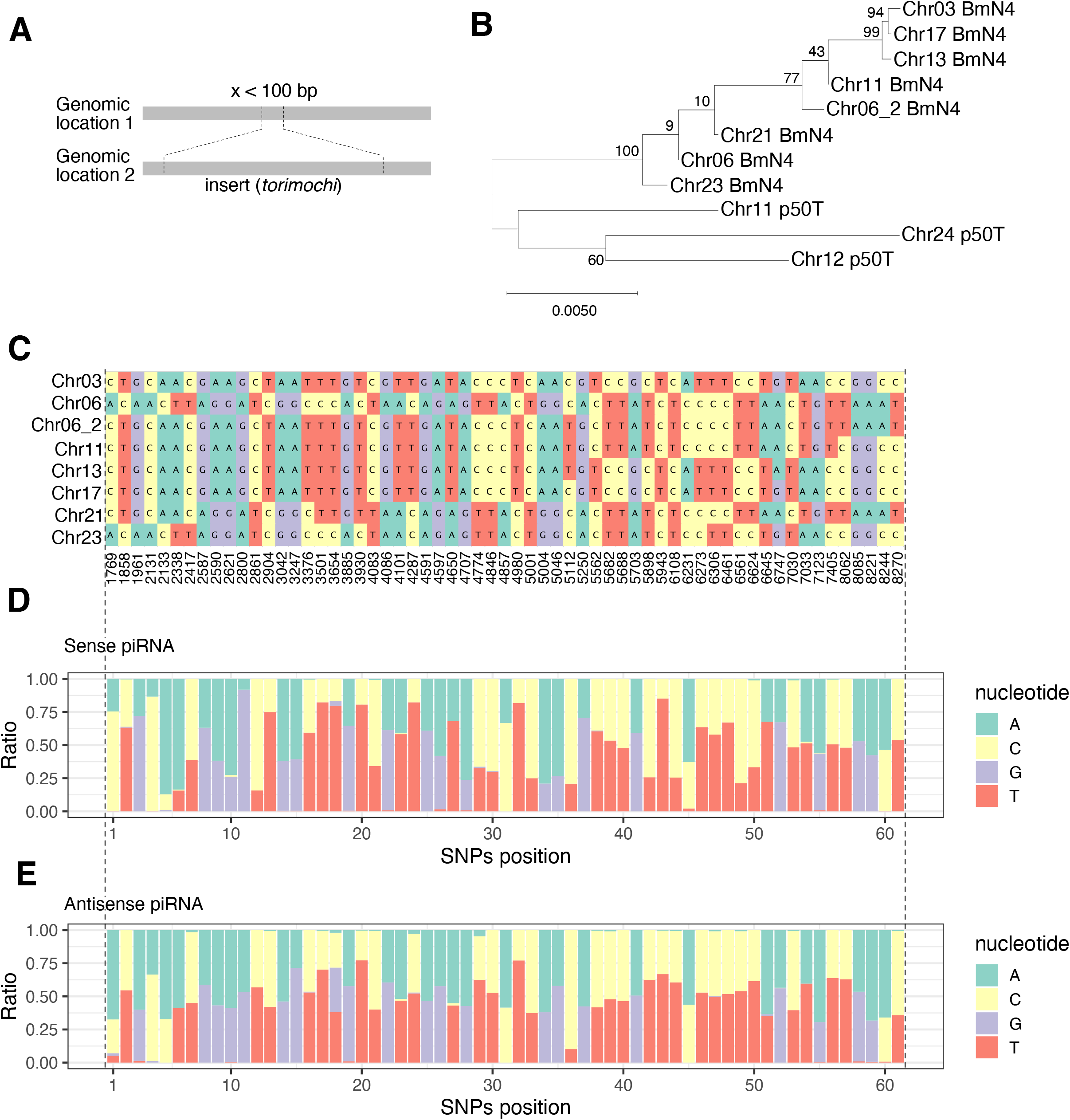
Detection of novel *torimochi* inserts in the BmN4 genome. (A) Definition of inserts in this paper. When the junctions between the two genomic locations are less than 100 bp apart at the genomic location 1 and are consistent in orientation, the region between the junctions at the genomic location 2 is considered as an insert. (B) enomic PCR using primers upstream and downstream of the newly identified *torimochi* inserts in BmN4 cells. Black and white arrows indicate the band lengths with and without *torimochi*, respectively. Chromosome 6 has at least two *torimochi* insertions: Chr06 and Chr06_2. (C) Phylogenetic tree of the new *torimochi* insert sequences in BmN4 cells and the *torimochi* copies present in p50T strain. Numbers indicate bootstrap probabilities. (D) List of SNPs in each *torimochi* in BmN4 cell. The numbers below indicate the position of the SNPs in *torimochi* on chromosome 3. (E, F) Distribution of SNPs at each position in sense (E) and antisense (F) piRNAs deriving from the new *torimochi* inserts in BmN4 cells. SNPs position indicates the order of the SNPs shown in (D).

In the previous *torimochi* paper, a GFP transgene, introduced via the piggyBac transposon, was trapped by *torimochi* and silenced by *de novo* piRNAs in BmN4 cells (Kawaoka et al. 2012). To identify which *torimochi* copy newly annotated in the BmN4 genome the GFP transgene was actually inserted into, we analyzed the genome of #8 cells, which produce abundant GFP-derived piRNAs (Kawaoka et al. 2012), using MinION.

We found two reads that cover the GFP cassette, *torimochi*, and the neighboring silkworm genome at the 11,929 kb position on chromosome 13 (Fig. S2I). Mapping of piRNAs showed that piRNA are produced mainly within the *torimochi* cassette rather than the surrounding regions (Fig. S2J), just like other *torimochi* copies on different chromosomes (Fig. S2A–H). These data suggest that, unlike the original assumption that *torimochi* is a unique, specialized piRNA cluster, the *torimochi* unit that captured the GFP transgene in #8 cells is merely one of many *torimochi* copies in the BmN4 genome.

### Comprehensive identification of transposons activated in the BmN4 genome

Given that *torimochi* turned out to be an active, multi-copy transposon, we decided to extend our analysis to other transposons in silkworms. We searched for novel transpositions in the BmN4 genome in an unbiased manner, using MinION sequencing and the same criteria as for *torimochi* (Fig. 2A). In this analysis, we defined “novel transpositions” when those sequences that existed in the p50T genome (published in 2019) are found elsewhere in the BmN4 genome. We identified ∼700 new inserts that have reliable junctions at both ends. The estimated size of these inserts was mostly in the range of 10^2^ to 10^4^ base pairs, so we decided to focus on the 597 inserts that fall within this range (Fig. 3A). These predicted inserts can be either full-length transposons, fragments of transposons, or transposon-unrelated sequences. Therefore, we categorized them into groups by sequence homology; when locally homologous sequences were bridged by a longer insert(s) (e.g., sequence 4), we considered them all as one group (Fig. 3B). Then, we further focused on 20 groups that include at least five inserts (Fig. 3C). Of these, remarkably, Group 2 that had the second most insert sites represented *torimochi* (and its fragments), while Group 1 with the highest number of inserts turned out to represent LINE transposon (and fragments). For each group, the longest sequence was selected as the representative and annotated by nucleotide BLAST with the *B. mori* transposon database (Table S2). As a result, they were identified to be LINE or SINE (Groups 1, 7, 9, 11, 15), known transposons (Groups 5, 6, 10, 12, 13, 16, 18, 19), *torimochi* (Group 2), or novel sequences (Groups 3, 4, 8, 14, 17, 20) (Table S2). We found that LTR transposons like *torimochi* and transposons with Inverted Terminal Repeat (ITR) tend to retain their full-length copies upon new transposition (Fig. 3D; Fig. S3E, G, H, I, K and L), whereas non-LTR transposons have produced incomplete copies fragmented from the 3’ side (Fig. S3A, B and C). These results suggests that the new insert sequences identified in this study are “alive” and active transposons that have transposed relatively recently. For these novel transposons, we named group 3 as *mejiro* (Japanese white-eye, which is often trapped by birdlime [torimochi]; see below), group 4 as *TRAS-lbm* (***TRAS-l****ike in* ***Bm****N4*; homologous to *TRAS* transposon), non-autonomous (transposase-lacking) transposons of group 8 as *kotaro*, 14 as *kojiro*, 17 as *kotetsu* (popular Japanese cat names; cats jump up and down, attracted by catnip [transposase from another autonomous transposon]), and group 20 *wao* (**w**andering p**ao**; homologous to *pao* transposon), respectively (Fig. S3A, B, E, G, I and L).

**Figure 3.**
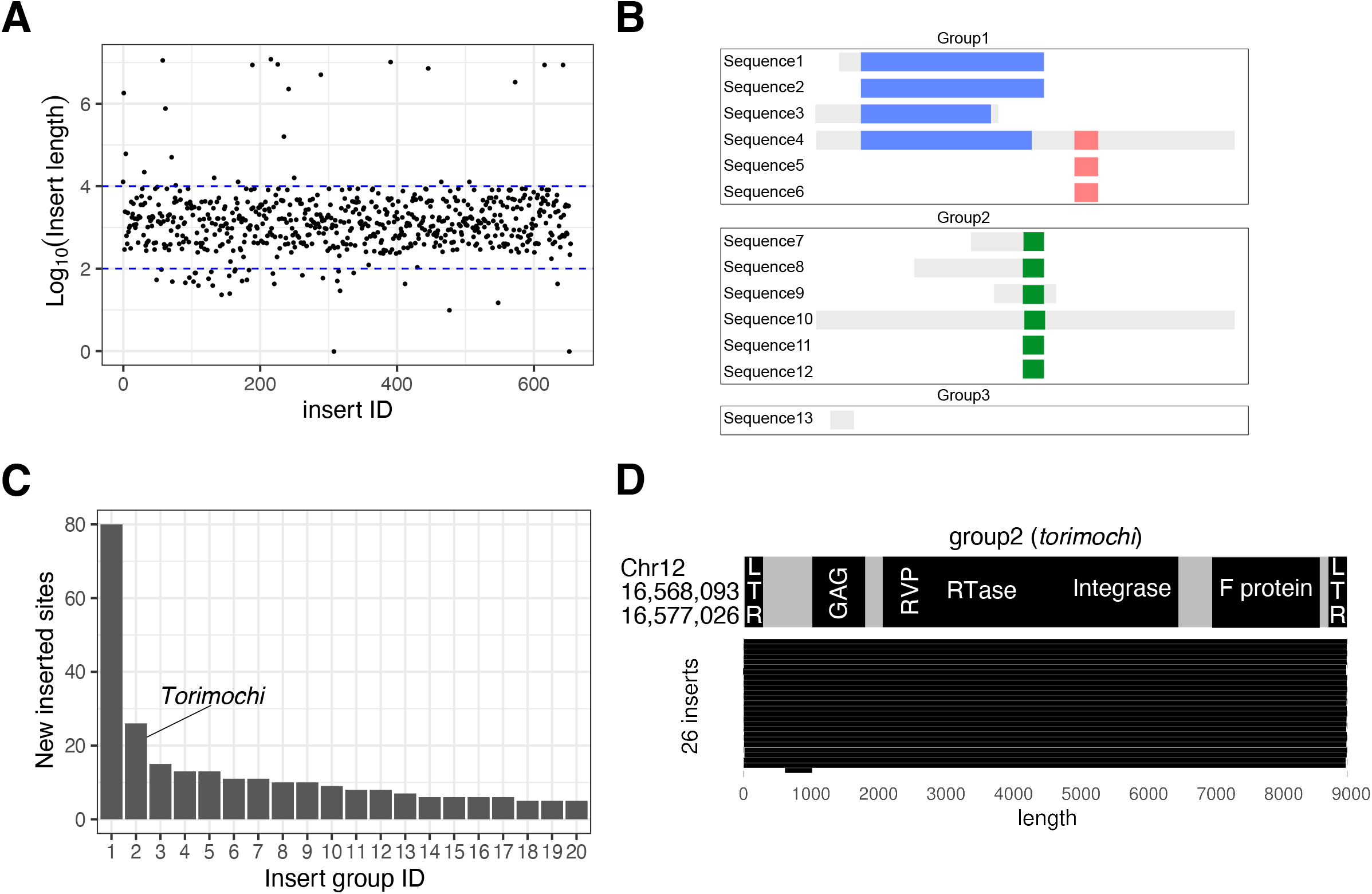
Comprehensive identification and grouping of novel insertion sequences. (A) Estimated lengths of novel inserts in the BmN4 genome. Blue dotted lines indicate 100 bp and 10,000 bp. (B) Schematic diagram of grouping. Segments with the same color are homologous to each other. For example, sequences 1 and 6 do not have direct homology but are bridged by sequence 4, thereby categorized into the same group. (C) The numbers of inserts in each group. Those groups that had 5 or more insert sites were sorted. Group 2 represents *torimochi*. (D) Domain structure of the representative sequence of group2 (*torimochi*) (top) and the region where it is originally annotated in the p50T genome (left). The lengths of the inserts found in BmN4 cells are shown at the bottom.

### *torimochi* is the most specifically activated transposon in BmN4 cells

Next, we compared the published genome sequence of the p50T strain and our MinION-based genome sequence of BmN4 cells. We found that p50T already has >70,000 copies of Group 1 (LINE-associated) insertions but only 10 copies of Group 2 insertions (including three full-length *torimochi* and seven partial *torimochi*; Table S3). To quantitatively evaluate the expansion of insertions in BmN4 cells, we calculated the ratio of the total genomic areas between the “old” copies already existing in the p50T genome and the “new” copies specifically found in the BmN4 genome for each insertion group. Notably, the ratio was the highest for *torimochi*, indicating that *torimochi* is the most massively expanded insertion in BmN4 cells (Fig. 4A). At the same time, we found that not only *torimochi* but also many other insertions (Groups 3, 8, 17, 4, 19 and 20) have increased their copies in BmN4 cells, most of which correspond to the novel transposons identified in this study (Fig. 4A).

**Figure 4.**
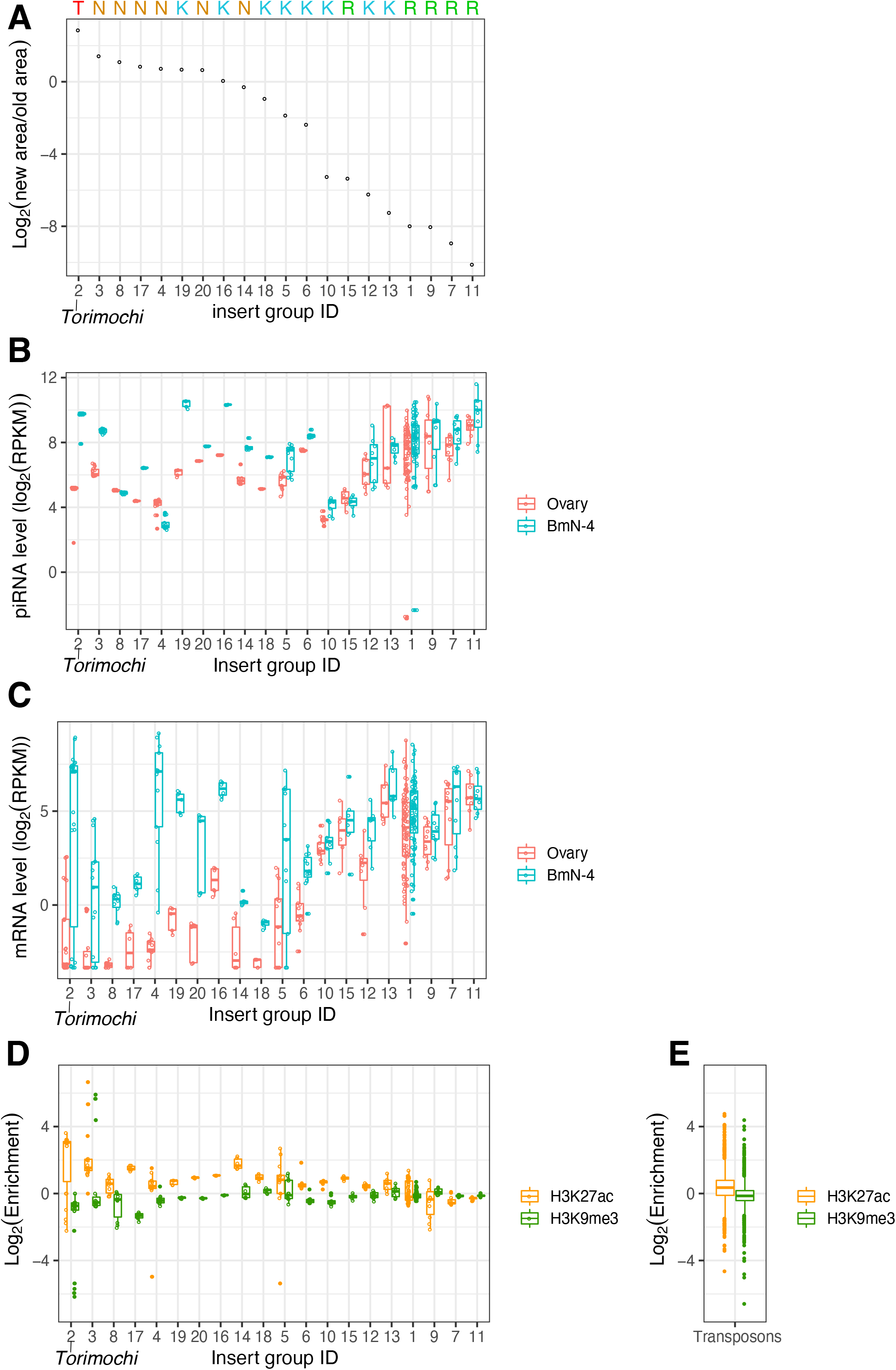
*torimochi* is the most active transposon in BmN4 cells. (A) Ratios between “new” areas in BmN4 cells and “old” areas in p50T strain for the inserts in each group. *torimochi* (group 2) is the most highly activated insert group in BmN4 cells. T (*torimochi*), N (novel transposons), K (known transposon), R (SINE/LINE) are shown (top). (B) The amounts of piRNAs derived from the inserts in each group in ovaries and BmN4 cells. The order is the same as in (A). Active inserts tend to produce more piRNAs in BmN4 cells than in ovaries. (C) The amounts of mRNAs derived from the inserts in each group in ovaries and BmN4 cells. The order is the same as in (A). Active inserts tend to produce more mRNAs in BmN4 cells than in ovaries. (D) Enrichment of the H3K27ac euchromatin mark and the H3K9me3 heterochromatin mark of the inserts in each group in ovaries and BmN4 cells. The order is the same as in (A). (E) Enrichment of the H3K27ac euchromatin mark and the H3K9me3 heterochromatin mark for 1811 known transposon sequences.

To investigate if these BmN4-specific new insertions have the ability to produce piRNAs like *torimochi*, we compared piRNA production from these insertions in silkworm ovaries and BmN4 cells (Fig. 4B). We found that most of the “new” insertions (left) show significantly increased piRNA production specifically in BmN4 cells, whereas the piRNA levels from “old” insertions such as LINE transposon (right) were comparable between silkworm ovaries and BmN4 cells (Fig. 4B). This increase of piRNA levels correlated well with the increase of mRNA levels in BmN4 cells (Fig. 4C). Previous ChIP-seq analysis revealed that *torimochi* is a transcriptionally active region with high levels of euchromatic modifications (Kawaoka et al. 2013). We therefore investigated the degree of histone modifications for these “new” inserts. Remarkably, these “new” insertions with piRNA-producing ability showed enrichment of the euchromatin mark H3K27ac and the depletion of the heterochromatin mark H3K9me3 (Fig. 4D), compared to their average levels in all sequences in the silkworm transposon database (Fig. 4E). These results suggest that these “new” insertions have an open chromatin structure, as previously reported for *torimochi* (Kawaoka et al. 2013, 2012). Nevertheless, it is noteworthy that, among those “new” insertions, *torimochi* has the highest activity in transposition and the most open chromatin structure (Fig. 4A and D), which may explain why the GFP transgene was trapped by *torimochi* for piRNA production in the previous study (Kawaoka et al. 2012). This is not limited to the GFP transgene; we found that the Group 3 transposon *mejiro*, the second most active transposon in BmN4 cells, is inserted into *torimochi* in a nested manner specifically in BmN4 cells (Fig. S4). Thus, *torimochi* is active in trapping not only exogenously introduced transgenes but also endogenous transposons.

## DISCUSSION

In this study, we found that *torimochi* is not a specialized piRNA cluster but in fact a full-length gypsy-type transposon that is exceptionally active in BmN4 cells. It was recently reported that large piRNA clusters are evolutionarily labile and can be deleted without compromising transposon regulation in *Drosophila* (Gebert et al. 2021). Thus, piRNA clusters in the genome are not as static as previously thought but can flexibly appear and disappear. Indeed, *torimochi* does not serve as a source of piRNAs in silkworm ovaries but has massively expanded its copy number and gained the activity to produce piRNAs in BmN4 cells (Fig. 1). *Torimochi* has the open chromatin structure and can trap foreign transgenes as well as endogenous transposons (Fig. 4, S2I and S4). Moreover, our SNP analysis showed that piRNAs are produced from many different copies of *torimochi* in the BmN4 genome. Therefore, *torimochi* in BmN4 cells may represent a young, growing piRNA cluster, which is still “alive” and active in transposition but capable of trapping other transposable elements to produce *de novo* piRNAs. Moreover, we successfully identified six novel transposons that have expanded their copies specifically in BmN4 cells, just like *torimochi* (Fig. S3). Future studies should focus on how transposons gain the ability of piRNA production and what hallmark features active piRNA clusters have.

## METHODS

### Cell lines

BmN4 cells (provided by Chisa Yasunaga-Aoki, Kyushu University, and maintained in our laboratory) were cultured at 27°C in IPL-41 medium (Applichem) supplemented with 10% fetal bovine serum.

### Search for *torimochi* sequences in the genome and their annotation

*torimochi* sequences were BLAST-searched in SilkBase (http://silkbase.ab.a.u-tokyo.ac.jp/) using the sequence information described in a previous paper (Kawaoka et al. 2012) as a query in the updated silkworm genome (Kawamoto et al. 2019). Since there were homologous regions on multiple chromosomes, the sequences were extended to the 5′ and 3′ ends, and then the common regions and Long Terminal Repeat (LTR) regions were identified by BLAST. The obtained nucleotide sequences were analyzed by pfam to estimate the domain structure of the encoded proteins (http://pfam.xfam.org/).

### Sequence analysis of deposited small RNAs

Small RNA sequence of BmN4 cells (DRR181866) and silkworm ovaries (DRR086591) were reported previously (Izumi et al. 2020; Katsuma et al. 2021). Informatics analysis of small RNAs was performed as reported previously (Shoji et al. 2017). In brief, the 3′-adaptor sequences were identified and removed, allowing for up to two mismatches. Reads of 20–42 nt were then obtained by excluding the reads shorter than 20 nt or longer than 42 nt. Small RNAs were mapped to the updated *Bombyx* genome (Kawamoto et al. 2019) using bowtie (Langmead et al. 2009). Reads that could be aligned to the genome up to two mismatches were used to calculate the mapping rate and to normalize each library. Sam files were converted to bam files by SAMtools (Li et al. 2009b), then to bed files, and the coverage of each nucleotide was calculated by BEDTools (Quinlan and Hall 2010).

### Genome extraction and genomic PCR

Genomic DNA was extracted from BmN4 cells using the Blood and Tissue kit (QIAGEN) according to the protocol of the manufacturer. Genomic DNA from *Bombyx mori* and *Bombyx mandarina* was provided by Susumu Katsuma. PCR was performed by step-down PCR using KOD One (TOYOBO), and the PCR products were analyzed in 1% agarose with 1 kb DNA Ladder (New England Biolabs). Primers used in this study are shown in Supplemental Table S1.

### Quantitative PCR for measuring the copy number of *torimochi*

Genomic DNA was used as a template for quantitative PCR analysis on a Thermal Cycler Dice Real Time System (TaKaRa) using KAPA SYBR Fast qPCR Kit (KAPA Biosystems) and the primers shown in Table S1 Absolute quantification of the copy number was performed using a known amount of plasmid containing the primer region. The primer set corresponding to the *rp49* gene on the autosome was used as a control.

### Cloning of *torimochi* and phylogenetic tree analysis

The sequences outside each *torimochi* copy were used to design the primers to distinguish it from other copies. PCR was performed by using inside-outside and outside-inside primer sets for each *torimochi* copy, and the PCR products were purified using NucleoSpin Gel and PCR Clean-up (MACHEREY-NAGEL) and then sequenced. *torimochi* sequences were clustered by Clustal W in MEGA X (Kumar et al. 2018) and the phylogenetic trees were generated by the maximum likelihood method with bootstrapping.

### SNPs detection

The sequence of each *torimochi* copy was combined and aligned with MEGA X (Kumar et al. 2018), and SNP-sites (Page et al. 2016) was used to generate a list of polymorphisms. The piRNA sequence library from previous literature (Izumi et al. 2020) was mapped to the *torimochi* sequence on chromosome 3 using bowtie (Langmead et al. 2009). Sam files were converted to bam files by SAMtools (Li et al. 2009b), and the SNP information of piRNAs was extracted using a custom R script.

### MinION-seq and mutation detection

Long DNA was extracted from BmN4 cells using Blood & Cell Culture DNA Kit (QIAGEN), sequencing libraries were prepared using Rapid Sequencing Kit (Oxford Nanopore), and the data was acquired by MinION using flow cell (version 9). The obtained sequences were mapped to the silkworm genome by ngmlr, and variants of genomic structure were detected by svim (Sedlazeck et al. 2018; Heller and Vingron 2019). Then, the structural variants of BND (breakends; connection of distal genomic positions) were extracted, and the inserts that have reliable junctions at both ends were obtained by using getfasta in BEDtools (Quinlan and Hall 2010).

### Mapping of ChIP-seq, mRNA-seq, and piRNA-seq to novel inserts and transposons

Previously published libraries of ChIP-seq, mRNA-seq and piRNA-seq were mapped to the newly identified inserts using hisat2 (ChIP-seq, mRNA-seq) and bowtie (piRNA-seq), and the numbers of length-corrected mapped reads were calculated (Langmead et al. 2009; Kim et al. 2019). Sam files were converted to bam files by SAMtools (Li et al. 2009b), then to bed files, and the coverage of each nucleotide was calculated by BEDTools (Quinlan and Hall 2010).

## DATA ACCESS

The sequencing data obtained in this study are available under the accession number DRA014297 (long DNA-seq). Small RNA sequences of BmN4 cells (DRR181866) and silkworm ovary (DRR086591), RNA sequences of BmN4 cells (DRR178519) and silkworm ovary (DRR186503), and ChIP sequences of IgG-R (DRR001873), H3K9me3 (DRR001878) and H3K27ac (DRR018643) antibodies were reported previously (Yokoi et al. 2021; Izumi et al. 2020; Katsuma et al. 2021, 2019; Kawaoka et al. 2013; Shoji et al. 2015).

## COMPETING INTEREST STATEMENT

The authors declare no competing interests.

## ACKNOWLEDGMENTS

We thank Susumu Katsuma and Takashi Kiuchi for providing silkworm genomic DNA and Kaori Kiyokawa for technical assistance. We thank all the members of the Tomari laboratory for discussion and critical comments on the manuscript. This work was supported in part by Grant-in-Aid for Scientific Research (S) (18H05271 to Y.T.), Grant-in-Aid for Young Scientists (Start-up) (19K06484 to K.S.)

## Author contributions

K.S., and Y.T. designed the study. K.S. and Y.U. performed molecular biological experiments. K.S. performed bioinformatic analysis of sequencing data. K.S. and Y.T. wrote the manuscript.

